# Lytic bacteriophage vB_KmiS-Kmi2C disrupts biofilms formed by members of the *Klebsiella oxytoca* complex, and represents a novel virus family and genus

**DOI:** 10.1101/2023.01.12.523727

**Authors:** Fiona Newberry, Preetha Shibu, Thomas Smith-Zaitlik, Mohamed Eladawy, Anne L. McCartney, Lesley Hoyles, David Negus

## Abstract

**AIMS:** This study aimed to characterise the lytic phage vB_KmiS-Kmi2C, isolated from sewage water on a GES-positive strain of *Klebsiella michiganensis*.

**METHODS AND RESULTS:** Comparative phylogenetic and network-based analyses were used to characterise the genome of phage vB_KmiS-Kmi2C (circular genome of 42,234 bp predicted to encode 55 genes), demonstrating it shared little similarity with other known phages. The phage was lytic on clinical strains of *K. oxytoca* (n=2) and *K. michiganensis* (n=4), and was found to both prevent biofilm formation and disrupt established biofilms produced by these strains.

**CONCLUSIONS:** We have identified a phage capable of killing clinically relevant members of the *Klebsiella oxytoca* complex (KoC). The phage represents a novel virus family (proposed name *Dilsviridae*) and genus (proposed name *Dilsvirus*).

**SIGNIFICANCE AND IMPACT OF THE STUDY:** Identification a novel lytic phage active against clinically relevant strains of the KoC provides an alternative to antibiotics to treat these increasingly antimicrobial-resistant opportunistic pathogens. The unusual way in which the phage can disrupt established biofilms may allow us to identify novel phage-based approaches for biofilm remediation in the future.

## INTRODUCTION

Bacteriophages (phages) are viruses of bacteria that infect their bacterial hosts for the purpose of replication. In the case of lytic phages, this process results in lysis and destruction of the host. The potential for applying the bactericidal action of lytic phages in a therapeutic setting was realised shortly after their discovery, with d’Herelle performing the first recorded clinical studies in 1918 (d’Herelle, 1918). Although interest in phage therapy diminished in Western countries following the introduction of antibiotics, the emergence of multidrug-resistant pathogens has resulted in a resurgence of interest.

Members of the *Klebsiella oxytoca* complex (KoC) are emerging as a serious health concern. The complex currently consists of several distinct phylogroups, representing *K. oxytoca* (Ko2), *K. michiganensis* (Ko1, Ko5), *K. grimontii* (Ko6), *K. huaxiensis* (Ko8), *K. pasteurii* (Ko4), *K. spallanzanii* (Ko3), and three unnamed novel species (Merla *et al*., 2019; Yang *et al*., 2022). Members are phylotyped based on the sequence of their chromosomally encoded β-lactamase (*bla*_oxy_) gene (Cosic *et al*., 2021). KoC bacteria can cause a range of serious infections in both humans and animals and are of particular concern in healthcare settings where immunocompromised patients are predisposed to potential infection. Isolates have been associated with urinary tract infections, lower respiratory tract infections, septicaemia, and soft-tissue injuries (Sohn, Seo & Jung, 2012; Paasch, Wilczek & Strik, 2017; Shakya *et al*., 2017; Lee *et al*., 2019). Certain members of the complex (*K. oxytoca, K. pasteurii, K. grimontii, K. michiganensis*) also encode the kleboxymycin biosynthetic gene cluster associated with antibiotic-associated haemorrhagic colitis (Shibu *et al*., 2021). Worryingly, antibiotic resistance amongst KoC members is increasing and resistance mechanisms include extended-spectrum beta-lactamases and carbapenemases (Lowe *et al*., 2012; White *et al*., 2016).

New treatment options for tackling drug-resistant infections caused by KoC isolates would be a welcome addition to our therapeutic armoury. This study aimed to characterise the morphology, genome and host range of a lytic phage, vB_KmiS-Kmi2C, isolated against a GES-5-encoding strain of *K. michiganensis* (PS_Koxy2) (Shibu *et al*., 2021). The anti-biofilm properties of the phage were investigated to further determine therapeutic utility with respect to preventing or resolving biofouling by KoC bacteria such as that which occurs on medical devices, particularly indwelling catheters and ventilators. Further extensive genomic and phylogenetic analyses of vB_KmiS-Kmi2C reveal that it represents a novel genus of phage.

## MATERIALS AND METHODS

### Strain and cultivation information

Details of all strains included in this study can be found in **Table 1**. All strains were grown on nutrient agar (Sigma Aldrich) unless stated otherwise. Nutrient broth (Sigma Aldrich) was used for overnight cultures unless stated otherwise, incubated aerobically at 37 °C. All media used for phage assays were supplemented with CaCl_2_ and MgCl_2_ (both at final concentration of 0.5 mM). The study of anonymised clinical isolates provided by the Nottingham University Hospitals NHS Trust (NUH) Pathogen Bank was approved by NUH Research and Innovation (19MI001).

**Table 1.** *K. oxytoca* complex strains used in this study

| Strain * | Species | Source | Reference |
| --- | --- | --- | --- |
| PS_Koxy1 | *K. michiganensis* | Throat swab | (Shibu *et al.*, 2021) |
| PS_Koxy2 §# | *K. michiganensis* | Urine | (Shibu *et al.*, 2021) |
| PS_Koxy4 | *K. michiganensis* | Rectal swab | (Shibu *et al.*, 2021) |
| Ko13 § | *K. michiganensis* | Blood culture | (Smith-Zaitlik *et al.*, 2022) |
| Ko14 | *K. michiganensis* | Blood culture | (Smith-Zaitlik *et al.*, 2022) |
| Ko21 § | *K. michiganensis* | Perfusion fluid | (Smith-Zaitlik *et al.*, 2022) |
| Ko22 § | *K. michiganensis* | Perfusion fluid | (Smith-Zaitlik *et al.*, 2022) |
| Ko37 § | *K. oxytoca* | Blood culture | (Smith-Zaitlik *et al.*, 2022) |
| Ko43 § | *K. michiganensis* | Bile fluid | (Smith-Zaitlik *et al.*, 2022) |
| Ko53 § | *K. oxytoca* | Blood culture | (Smith-Zaitlik *et al.*, 2022) |

\* Ko prefix indicates strains came from the Pathogen Bank of Queen's Medical Centre Nottingham, UK.
§ Infected by phage vB\_KmiS-Kmi2C.

### Generation of hybrid genome sequences

Cell pellets from 10-ml overnight cultures grown in nutrient broth were sent to microbesNG (Birmingham, UK) for DNA extraction, library preparation and sequencing according to the sequencing provider’s strain-submission procedures (refer to https://microbesng.com/documents/24/MicrobesNG_Sequencing_Service_Methods_v20210419.pdf for the protocol).

For Illumina sequencing, the Nextera XT Library Preparation Kit (Illumina, San Diego, USA) was used and paired-end reads (HiSeq/NovaSeq; 2x250 bp were generated for PS_Koxy1 (coverage 141×), PS_Koxy2 (coverage 52×) and PS_Koxy4 (coverage 57×). Reads were adapter-trimmed using Trimmomatic 0.30 (Bolger, Lohse & Usadel, 2014) with a sliding window quality cut-off of Q15. For Illumina/MinION hybrid genomes, long-read genomic DNA libraries were prepared with Oxford Nanopore SQK-RBK004 kit and/or SQK-LSK109 kit with Native Barcoding EXP-NBD104/114 (ONT, United Kingdom) using 400–500 ng of high-molecular-weight DNA. Barcoded samples were pooled together into a single sequencing library and loaded into a FLO-MIN106 (R.9.4.1) flow cell in a GridION (ONT, United Kingdom). *De novo* genome assembly was performed using Unicycler v0.4.0 (Wick *et al*., 2017).

The genome assemblies returned to us by microbesNG were annotated in-house using Bakta v1.6.1, database v4.0 (Schwengers *et al*., 2021). RGI 6.0.0, CARD 3.2.5 was used to predict antimicrobial resistance genes encoded within genomes (Alcock *et al*., 2020). COPLA and plaSquid were used to characterise plasmids harboured by strains (Redondo-Salvo *et al*., 2021; Giménez, Ferrés & Iraola, 2022).

### Biolog GEN III MicroPlate assays

All strains were grown aerobically overnight on blood-free BUG agar (Biolog) at 37 °C. For each assay, one to two colonies were used to inoculate Biolog Inoculation Fluid B to a turbidity of 95 % (Biolog turbidimeter). Biolog Gen III plates were inoculated according to the manufacturer’s instructions, and incubated aerobically at 37 °C for 22 h. Results were read at 540 nm using a BioTek Cytation |^3^ imaging reader spectrophotometer. All Biolog reagents and kits were purchased from Technopath. Assays were carried out in triplicate for strains PS_Koxy1, PS_Koxy2 and PS_Koxy4.

### Biofilm assays and their interpretation

Biofilm assays were performed as described previously (Stepanovic *et al*., 2000; Merritt, Kadouri & O’Toole, 2005; Eladawy *et al*., 2021). In brief, a single colony of each strain was used to inoculate 5 ml of tryptone soy broth supplemented with 1 % glucose (TSBG). Cultures were incubated aerobically for 24 h at 37 °C without shaking. The overnight cultures were diluted to 1:100 using TSBG, then aliquots (100 µl) of the diluted cultures were introduced into wells of a 96-well plate. The plates were incubated aerobically for 24 h at 37 °C without shaking. Then, the spent medium was carefully removed from each well. The wells were washed three times with 200 µl sterile phosphate-buffered saline (pH 7.4; Oxoid) to remove any non-adherent planktonic cells. The adherent cells were fixed by heat treatment at 60 °C for 60 min to prevent widespread detachment of biofilms prior to dye staining. The adhered biofilms were then stained by addition of 1 % (w/v) crystal violet (150 µl per well) and the 96-well plate was left to incubate for 20 min. The excess stain was then carefully removed from the wells and discarded. The 96-well plate was then carefully rinsed with distilled water three times, and the plate was inverted and left at room temperature until the wells were dry. The stained biofilms were solubilised by adding 33 % (v/v) glacial acetic acid (Sigma Aldrich) to each well (150 µl per well). After solubilisation of the stained biofilms, the OD_540_ was measured and recorded for all samples using a BioTek Cytation |^3^ imaging reader spectrophotometer. *K. michiganensis* Ko14 and uninoculated medium were used as negative controls in biofilm assays. Biological (*n*=3) and technical (*n*=4) replicates were done for all strains. The mean of each isolate’s OD quadruplicate readings (OD_i_) was calculated and compared with the control cut-off OD (OD_C_), which was defined as three standard deviations (SD) above the mean of the negative control (3SD + mean). The amount of biofilm formed was scored as non-adherent (OD_i_ ≤ OD_C_), weakly adherent (OD_C_ < OD_i_ ≤ 2 OD_C_), moderately adherent (2 OD_C_ < OD_i_ ≤ 4 OD_C_) or strongly adherent (4 OD_C_ < OD_i_).

### Isolation of lytic phage and host range analysis

Filter-sterilised sewage samples (0.45 µm cellulose acetate filter; Millipore) collected from mixed-liquor tanks at Mogden Sewage Treatment Works (March 2017) were screened against *K. michiganensis* strain PS_Koxy2 as described previously (Smith-Zaitlik *et al*., 2022). Host range analysis was carried out as described previously (Smith-Zaitlik *et al*., 2022). After overnight incubation, overlay assay plates were inspected for lysis, with results recorded according to a modification of (Haines *et al*., 2021): ++, complete lysis; +, hazy lysis; 0, no visible plaques.

### Phage concentration and transmission electron microscopy (TEM)

The Vivaspin 20 50 kDa centrifugal concentrator (Cytiva) was used to concentrate filter-sterilised propagated phage as described previously (Smith-Zaitlik *et al*., 2022). For TEM, formvar/carbon-coated 200 mesh copper grids (Agar Scientific) were prepared via glow discharge (10 mA, 10 s) using a Q150R ES sputter coater (Quorum Technologies Ltd) and processed as described previously (Smith-Zaitlik *et al*., 2022). Samples were visualised using a JEOL JEM-2100Plus (JEOL Ltd) TEM and an accelerating voltage of 200 kV. Images were analysed and annotated using ImageJ (https://imagej.net/Fiji).

### Phage DNA extraction and sequencing

Phage DNA was extracted from concentrated phage lysate using the Qiagen DNeasy Blood & Tissue Kit as described previously (Smith-Zaitlik *et al*., 2022). Sequence data were generated on our in-house Illumina MiSeq platform, with the Nextera XT DNA library preparation kit (Illumina) to produce fragments of approximately 500 bp. Fragmented and indexed samples were run on the sequencer using a Micro flow cell with the MiSeq Reagent Kit v2 (Illumina; 150-bp paired-end reads) following Illumina’s recommended denaturation and loading procedures.

### Phage genome assembly and annotation

Quality of raw sequence data was assessed using <u>FastQC</u> v0.11.9. Reads had a mean phred score above 30 and contained no adapter sequences, so data were not trimmed. The genome of the novel phage was assembled using SPAdes v3.15.4 (default settings) (Bankevich *et al*., 2012) and visualised using Bandage v0.8.1 (Wick *et al*., 2015). Contamination and completeness of the genome was determined using CheckV v0.8.1 (CheckV database v1.0) (Nayfach *et al*., 2021). Gene annotations were made using Prokka (v1.14.6) with the PHROGs database (v3) (Terzian *et al*., 2021). The genome was screened for antimicrobial resistance genes using the Resistance Gene Identifier (v5.2.0) of the Comprehensive Antibiotic Resistance Database (CARD) (v3.1.4) (Alcock *et al*., 2020), and for virulence genes using the Virulence Factor Database (VFDB) (accessed: 6/6/2022) (Liu *et al*., 2019). Presence of temperate-lifestyle genes was determined using PhageLeads (Yukgehnaish *et al*., 2022). An annotated genome map was generated using GenoPlotR (v0.8.11) and predicted protein function from the PHROGs database (Guy, Kultima & Andersson, 2010). To identify potential depolymerase-associated genes encoded within the phage genome, predicted protein names were searched for the following terms: ‘depolymerase’, ‘pectin’, ‘pectate’, ‘sialidase’, ‘levanase’, ‘xylosidase’, ‘rhamnosidase’, ‘dextranase’, ‘alginate’, ‘hyaluronidase’, ‘hydrolase’, ‘lyase’.

### Comparative genome analysis

To generate a gene-sharing network, 21,903 phage genomes from the INPHARED database (April 2022) and the query phage were clustered using vConTACT2 (v0.9.22; default settings) and visualised in Cytoscape (v.3.9.1) (Bolduc *et al*., 2017; Cook *et al*., 2021). The viral cluster (VC) containing the query phage was determined and a proteomic phylogenetic tree generated from all phage within the same VC using VipTree (v1.1.2; default settings) (Nishimura *et al*., 2017). *Klebsiella oxytoca* phage genomes vB_KmiM-2Dii (accession: MZ707156) and vB_KmiM-4Dii (accession: MZ707157) were used as outliers. An additional proteomic phylogenetic tree was generated from all phages within the VC and three representative phage genomes from distantly related genera *Nonagvirus* (JenK1; NC_029021.1), *Seuratvirus* (CaJan; NC_028776.1) and *Nipunavirus* (NP1; NC_031058.1). The large terminase protein sequences were used to generate a maximum likelihood phylogenetic tree with ClustalW (v.2.1) and IQTree with 1000 bootstraps (v1.6.12) (Thompson, Higgins & Gibson, 1994; Minh, Nguyen & von Haeseler, 2013). IQModelFinder (v.1.4.2) was used with IQTree to determine the model of best fit (Kalyaanamoorthy *et al*., 2017). The percentage of shared proteins within the VC was determined using all-versus-all BLASTP (≥30 % identity and ≥50 % sequence coverage; v2.12.0) (Turner, Kropinski & Adriaenssens, 2021). The intergenomic similarity between phage within the same VC was determined using VIRIDIC (v1.0) (Moraru, Varsani & Kropinski, 2020). ComplexHeatmap was used to generated all heatmaps (Gu, Eils & Schlesner, 2016).

### Phage–biofilm assays

The titre of the phage stock was determined by plaque assay using the double-layer agar technique. Briefly, phage vB_KmiS-Kmi2C was serially diluted in phosphate-buffered saline (pH 7.4; Oxoid) and 100 µl of each phage dilution was combined with 100 µl of an overnight culture of *K. michiganensis* PS_Koxy2 and 5 ml of 0.6 % tryptone soy agar (TSA) supplemented with CaCl_2_ and MgCl_2_ both at a final concentration of 1 mM. The mixture was gently swirled and poured onto solid TSA plates. Plates were incubated overnight at 37 °C and pfu/ml determined by enumeration of visible plaques.

The ability of vB_KmiS-Kmi2C to prevent and disrupt biofilms was examined using a modification of a previously described protocol (Taha *et al*., 2018). Hosts of vB_KmiS-Kmi2C identified as moderately (2 OD_C_ < OD_i_ ≤ 4 OD_C_) adherent were included in the assay. For prevention of biofilms, host cultures were incubated aerobically for 24 h at 37 °C without shaking in TSBG. Overnight cultures were diluted 1:100 with TSBG and aliquots (100 µl) of diluted culture were introduced into wells of a 96-well plate with or without phage vB_KmiS-Kmi2C (added to a final concentration of 4.5 x 10^8^ pfu/ml to each test well). Plates were incubated without shaking for 24 h at 37 °C. Then, the supernatants were discarded, the biofilm of each well was washed to remove planktonic cells and biofilms stained as described above.

To investigate the disruption of established biofilms, host cultures were grown and prepared as described above prior to inoculating a 96-well plate. Plates were incubated without shaking for 24 h at 37 °C to allow biofilms to form. Unattached planktonic cells were carefully aspirated without disrupting the biomass. Phage vB_KmiS-Kmi2C diluted in 100 µl TSBG was added to test wells (final titre of 4.5×10^8^ pfu/ml) whereas control wells received only TSBG without phage. Plates were incubated for a further 24 h at 37 °C without shaking. Supernatants were carefully discarded; the biofilm of each well was washed to remove planktonic cells and biofilms stained as described above. Biological (*n*=3) and technical (*n*=4) replicates were completed for all strains.

## RESULTS

### Host range of phage vB_KmiS-Kmi2C

Phage vB_KmiS-Kmi2C was isolated from sewage water on *K. michiganensis* PS_Koxy2 (ST138), a multidrug-resistant, GES-5-positive isolate recovered from human urine (Shibu *et al*., 2021). The phage was screened against 56 clinical and 28 animal isolates representing a range of *Klebsiella* spp. (*K. michiganensis* n=49, *K. oxytoca* n=25, *K. grimontii* n=9, *K. huaxiensis* n=1) (Shibu *et al*., 2021; Smith-Zaitlik *et al*., 2022). It did not infect the closely related *K. michiganensis* strains PS_Koxy1 and PS_Koxy4 (Shibu *et al*., 2021). vB_KmiS-Kmi2C did not infect animal isolates of *Klebsiella* spp. (Smith-Zaitlik *et al*., 2022), but did infect some of the clinical *K. oxytoca* (*n*=2) and *K. michiganensis* (*n*=4) strains within our extended in-house collection (Smith-Zaitlik *et al*., 2022) (**Table 1**). From strain information provided by the Pathogen Bank (EUCAST testing) all these strains were resistant to amoxycillin, and sensitive to amikacin, ceftazidime, ceftazidime, ceftriaxone, cefuroxime, ciprofloxacin, ertapenem, gentamicin, meropenem, piptazobactam and trimethoprim. Strains Ko13, Ko22, Ko43 and Ko53 were sensitive to co-amoxiclav; Ko21 was resistant to co-amoxiclav; no data were supplied for Ko37 and co-amoxiclav. Phage vB_KmiS-Kmi2C did not exhibit depolymerase activity (i.e. it did not form haloes around plaques) on any of the strains that it infected.

### Genotypic and phenotypic characterisation of the GES-positive strains

Our previous work had shown PS_Koxy1, PS_Koxy2 and PS_Koxy4 to be very similar based on the analysis of draft genome sequence data (Shibu *et al*., 2021). In this study, we generated hybrid assemblies of their genomes (**Table 2**).

**Table 2.** Summary statistics for hybrid genome assemblies of the GES-5-positive strains

| Strain (accession) | Contigs | Size (bp) | Status * | N50 | CDS | Plasmids (RIP; MOB; REP; PTU) # |
| --- | --- | --- | --- | --- | --- | --- |
| PS_Koxy1<br>(GCA_014050555.2) | 7 | 6,392,093 | CCC; 4 CCPs; ?<br>incomplete<br>plasmid | 6,023,296 | 5,952 | pPSKoxy1_1 (RepA_C; MOBH; IncAC, IncN; -)<br>pPSKoxy1_2 (Rep_3; -, IncR; -)<br>pPSKoxy1_3 [-; MOBP1; -, PTU-Q2 (IncQ)]<br>pPSKoxy1_4 (-; MOBP1; Col440I; PTU-E3) |
| PS_Koxy2<br>(GCA_014050535.2) | 10 | 6,357,486 | Draft<br>chromosome; 4<br>CCPs; ?<br>incomplete<br>plasmid | 5,917,750 | 5,910 | pPSKoxy2_1 (IncFII_repA, IncFII_repA, RepA_C, RepB-RCR_reg; MOBF, MOBH; IncFII_1_pKP91, IncAC, IncN; -)<br>pPSKoxy2_2 (Rep_3; -, IncR; -)<br>pPSKoxy2_3 [-; MOBP1; -, PTU-Q2 (IncQ)]<br>pPSKoxy2_4 (-; MOBP1; Col440I; PTU-E3) |
| PS_Koxy4<br>(GCA_014050515.2) | 6 | 6,406,207 | CCC; 5 CCPs | 6,075,855 | 5,969 | pPSKoxy4_1 [IncFII_repA, IncFII_repA, RepB-RCR_reg; MOBF; IncFII_1_pKP91; PTU-FK (IncFII)]<br>pPSKoxy4_2 [-; MOBH; IncAC; PTU-C (IncA)]<br>pPSKoxy4_3 (Rep_3; -, IncR; -)<br>pPSKoxy4_4 [-; MOBP1; -, PTU-Q2 (IncQ)]<br>pPSKoxy4_5 (-; MOBP1; Col440I; PTU-E3) |

\*CCC, complete circular chromosome; CCP, complete circular plasmid.

The initial description of the strains’ draft genomes (Ellington *et al*., 2019) predicted they each harboured an identical IncQ 8,300-bp circular GES-5-positive plasmid, confirmed in this study. PS_Koxy2 and PS_Koxy4 both carried an identical 76,870-bp circular plasmid (replication initiation protein domain Rep_3, replicon type IncR). The 4,448-bp plasmid carried by all three strains shared high identity with seven other plasmids, harboured by *Salmonella enterica*, *Enterobacter* spp. and *K. michiganensis* (**Table 3**). A comparison (progressiveMauve, not shown) of the sequence of pPSKoxy2_4 with those of the other 4,448-bp plasmids in **Table 3** showed they shared 4,446 identical sites (99.98 % pairwise identity) across their length. A PlasmidMLST search returned no hits for the plasmid sequence. It was found to be PTU-E3 (score 96.97 %) by COPLA, exceeding the 90 % threshold to validate the plasmid taxonomic unit assignment [16], and to belong to mobility group MOBP1 with a Col440I replicon type by plaSquid (Giménez *et al*., 2022). The plasmid was predicted by Bakta to encode six genes (MobA, MobC, two YgdI/YgdR family lipoproteins, two hypothetical proteins), an origin of replication, *oriT* and a non-coding RNAI (**Figure 1a**). UniProt BLASTP analyses showed MobA shared 55 % identity (across 517 aa) with the DNA relaxase MbeA of *Escherichia coli* (reviewed UniProt record P13658); MobC shared 63.6 % identity (across 115 aa) with the mobilisation protein MbeC of *E. coli* (reviewed UniProt record P13657). The 39-aa hypothetical protein was predicted by I-TASSER (Zhang, 2008) to be an alpha helix, while the 122-aa hypothetical protein was found to include a transmembrane segment (HHPRED, UniRef30 (Zimmermann *et al*., 2018)) that had limited structural homology with only the *Escherichia coli* signal transduction protein PmrD (PDB:4HN7) and the Vps26 dimer region of a fungal vacuolar protein sorting-associated protein (PDB:7BLQ).

**Figure 1.**
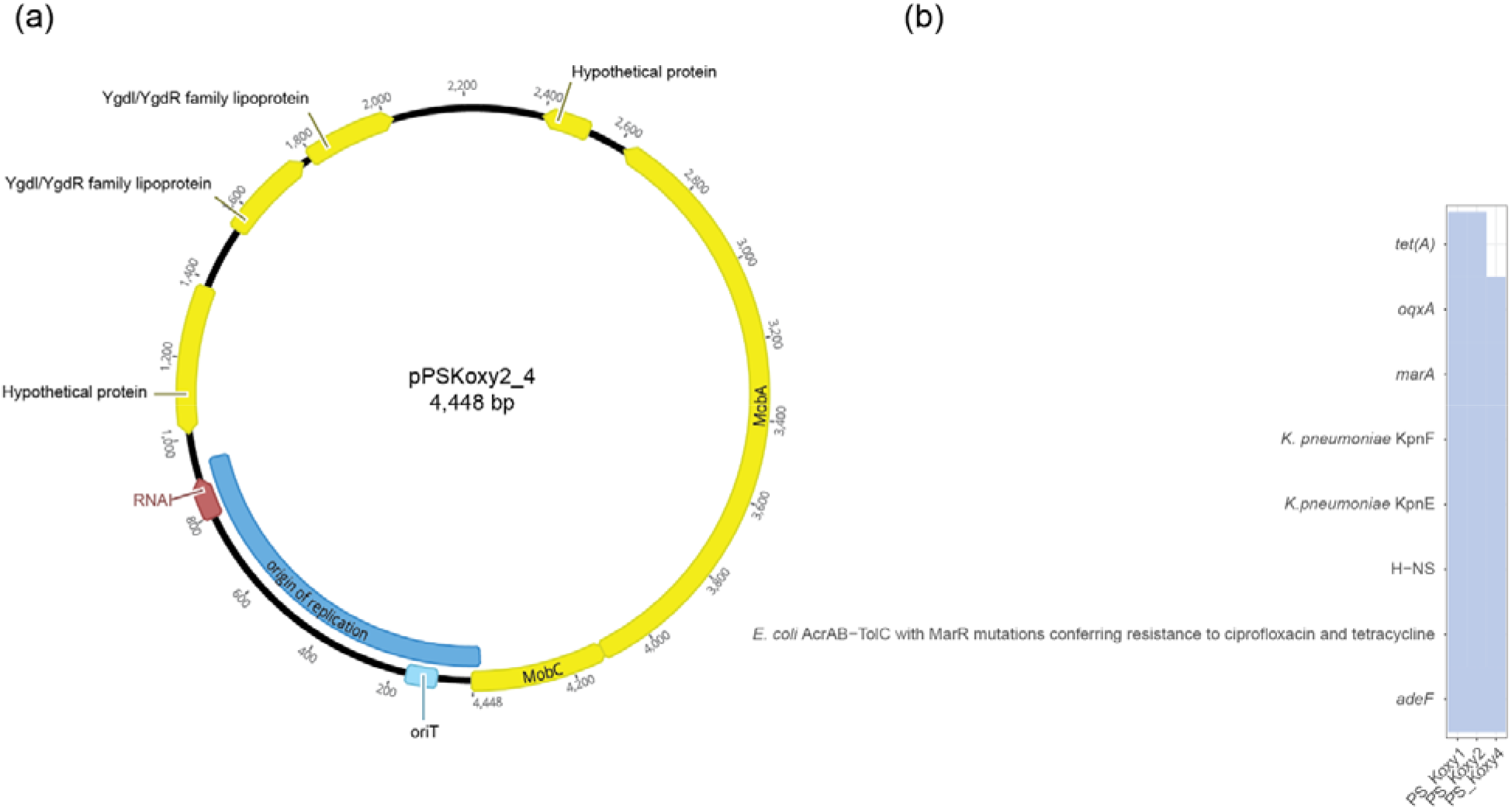
**(a)** Illustration of the circular conjugative 4,448-bp plasmid harboured by *K. michiganensis* strains PS_Koxy1, PS_Koxy2 and PS_Koxy4. **(b)** Tetracycline-antibiotic-associated genes predicted to be encoded within the genomes of strains PS_Koxy1, PS_Koxy2 and PS_Koxy4 by RGI/CARD. All hits were considered ‘strict’ by CARD.

**Table 3.** Closest relatives of the circular 4,448 bp plasmid as determined using NCBI BLASTN

| Species, strain and plasmid | Identity | Size (nt) | Accession | Source | Location | Reference |
| --- | --- | --- | --- | --- | --- | --- |
| *S. enterica* PNUSAS021403 pPNUSAS021403_3 | 3511/3513 (99 %) | 4,448 | CP093106.1 | Patient | USA | (Webb *et al.*, 2022) |
| *S. enterica* subsp. *enterica* serovar Hadar PNUSAS018090 pPNUSAS018090_3 | 2662/2664 (99 %) | 4,448 | CP093099.1 | Patient | USA | (Webb *et al.*, 2022) |
| *S. enterica* 2016K-0377 p2016K-0377_3 | 4387/4389 (99 %) | 4,448 | CP093079.1 | Patient | USA | (Webb *et al.*, 2022) |
| *S. enterica* subsp. *enterica* serovar Hadar 12-2388 p12-2388.3 | 2662/2664 (99 %) | 4,448 | CP038598.1 | Reference laboratory | Canada | (Yachison *et al.*, 2017) |
| *E. cloacae* 78 pP6 | 3403/3403 (100 %) | 9,923 | OW849463.1 | Hospital | Spain | BioProject PRJEB42440 |
| *E. hormaechei* RHBSTW-00070 pRHBSTW-00070_3 | 3403/3403 (100 %) | 9,923 | CP058166.1 | Wastewater influent | UK | (AbuOun *et al.*, 2021) |
| *K. michiganensis* RHBSTW-00676 pRHBSTW-00676_6 | 3211/3211 (100 %) | 4,448 | CP055990.1 | Wastewater influent | UK | (AbuOun *et al.*, 2021) |

PS_Koxy4 could be differentiated from PS_Koxy1 and PS_Koxy2 by phenotypic traits determined using the Biolog GEN III system (**Table 4**). PS_Koxy1 and PS_Koxy2 were resistant to the tetracycline antibiotic minocycline; both were predicted to encode the tetracycline efflux pump *tet(A)* on incompletely assembled plasmids (contig_3 PS_Koxy1, predicted to be FIIK by PlasmidMLST; contig_2 PS_Koxy2, predicted to be IncA/C ST12 by PlasmidMLST), but this gene was absent from the genome of PS_Koxy4, with the strain sensitive to minocycline (**Figure 1b**, **Table 4**).

**Table 4.** Phenotypic traits that differentiate PS_Koxy1 and PS_Koxy2 from PS_Koxy4

| Growth in presence of | PS_Koxy1 | PS_Koxy2 | PS_Koxy4 |
| --- | --- | --- | --- |
| D-Serine | – | – | + |
| L-Alanine | + | + | – |
| Minocycline | + | + | – |
| Acetic acid | – | – | + |

+, Positive according to the Biolog GEN III system; –, negative according to the Biolog GEN III system.

### Biofilm-forming ability of strains used in this study

We determined the biofilm-forming ability of strains used in this study as described previously (Taha *et al*., 2018). Our results show a wide range of biofilm-forming abilities among the strains (**Figure 2a**; **Supplementary Table 1**), yet there was no difference in the virulence factor profiles of PS_Koxy1, PS_Koxy2 and PS_Koxy4 according to a VFDB (Liu *et al*., 2019) analysis (**Supplementary Table 2**). Strains *K. michiganensis* PS_Koxy1, *K. michiganensis* Ko13, *K. michiganensis* Ko21 and *K. michiganensis* Ko22 were identified as weakly adherent (OD_C_ < OD_i_ ≤ 2 OD_C_). Strains *K. michiganensis* PS_Koxy2, *K. michiganensis* PS_Koxy4, *K. oxytoca* Ko37, *K. oxytoca* Ko43 and *K. oxytoca* Ko53 were identified as moderately adherent (2 OD_C_ < OD_i_ ≤ 4 OD_C_). *K. michiganensis* Ko14 was used as a non-adherent negative control (OD_i_ ≤ OD_C_).

**Figure 2.**
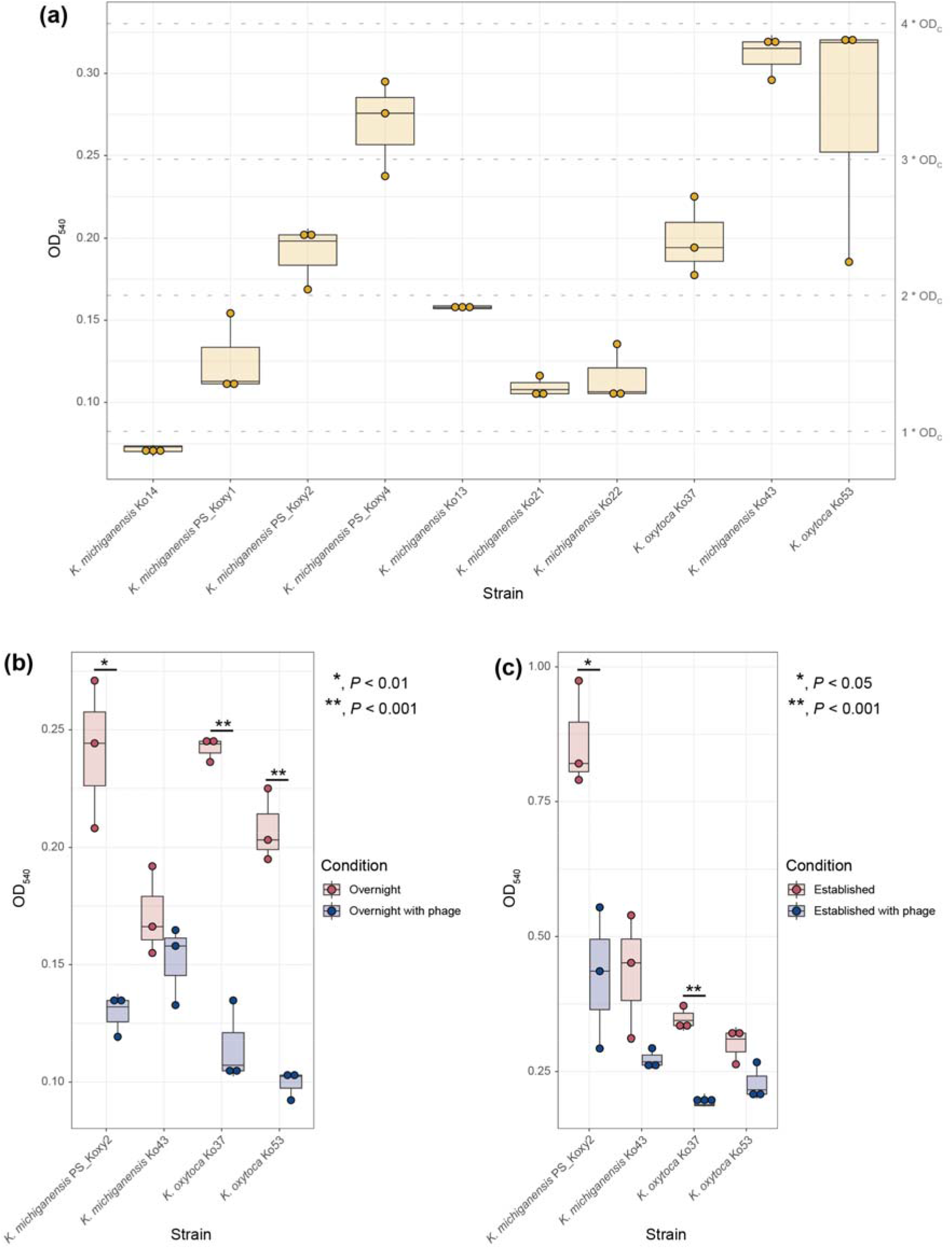
Biofilm-forming abilities of the strains used in this study. **(a)** The biofilm-forming abilities of all strains listed in **Table 1** were assessed. *K. michiganensis* Ko14, a strain that does not form biofilms in TSBG, was used as the negative control. Differences in biofilm formation in comparison with the control were statistically significant for all strains (unpaired *t* test, adjusted *P* value < 0.05, Benjamini-Hochberg; **Supplementary Table 1**). The ability of phage vB_KmiS-Kmi2C to **(b)** prevent biofilm formation or **(c)** disrupt established biofilms was assessed using those strains infected by the phage and forming the strongest biofilms **(a)**. **(b, c)** Statistical significance of the differences in biofilm formation of strains in the presence and absence of phage was assessed using unpaired *t* test. **(a–c)** Data are shown as mean for four technical and three biological replicates per strain.

### Capacity of phage vB_KmiS-Kmi2C to prevent and disrupt biofilms

Phage vB_KmiS-Kmi2C was found to be highly effective at both preventing (**Figure 2b**) and disrupting (**Figure 2c**) biofilms formed by isolates identified as susceptible to lysis. Presence of vB_KmiS-Kmi2C was found to reduce biofilm formation for all strains included in the study. This reduction was found to be significant (*P* < 0.01) for three of the isolates tested (PS_Koxy2, Ko37 and Ko53) compared to no-phage controls. Addition of vB_KmiS-Kmi2C to pre-established biofilms also resulted in biofilm disruption, seen as a reduction in measured biofilm compared to non-phage-treated controls. Biofilm disruption was determined to be significant (*P* < 0.05) for isolates PS_Koxy2 and Ko37 compared to non-phage treated controls.

### Morphology of phage vB_KmiS-Kmi2C

TEM showed vB_KmiS-Kmi2C to have a siphovirus-like morphology (**Figure 3**). It had a long, non-contractile tail. The capsid diameter was 59.6 nm (SD 2.3 nm), the tail and baseplate were 157.8 nm in length (SD 5.1 nm), and the phage had a total length of 218.11 nm (SD 4.8) (*n*=3 measurements).

**Figure 3.**
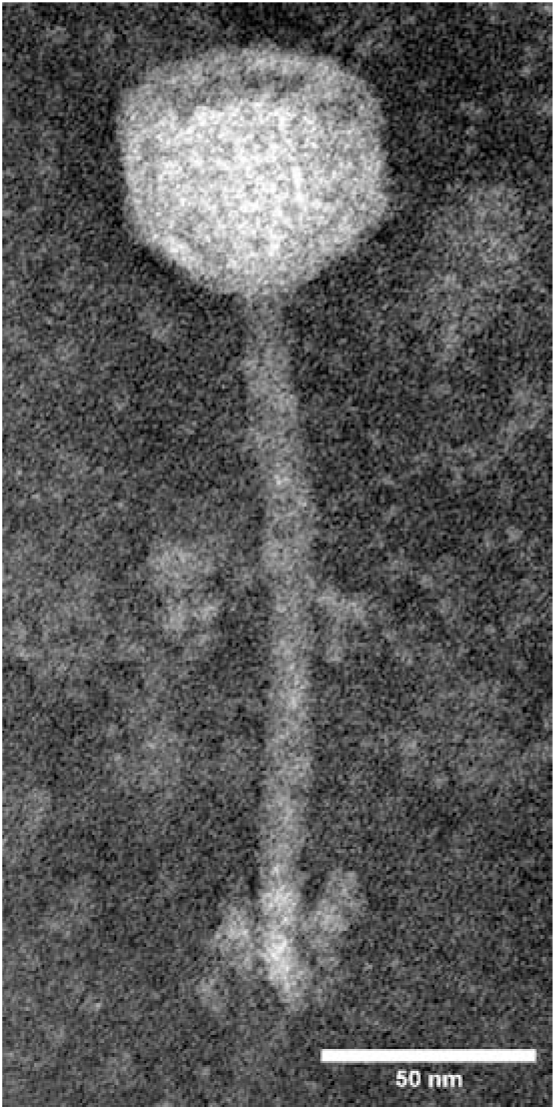
Transmission electron micrograph of phage vB_KmiS-Kmi2C. Scale bar, 50 nm.

### Genome of phage vB_KmiS-Kmi2C

The genome of phage vB_KmiS-Kmi2C was assembled into a single contig, with 100 % completeness according to CheckV (Nayfach *et al*., 2021). Assembly of the phage was confirmed using visualisation with Bandage (**Supplementary Figure 1**). It comprised 42,234 bp (816 × coverage) and was predicted to encode 55 genes (upper panel **Figure 4**). The genome is arranged into functional modules as typically seen in phage genomes: replication/regulation, viral structure, DNA packaging and lysis. Interesting genomic features of the genome included a Sak4-like ssDNA annealing protein with single-strand DNA-binding protein, a Cas4-domain exonuclease, a holin and a Rz-like spanin.

**Figure 4.**
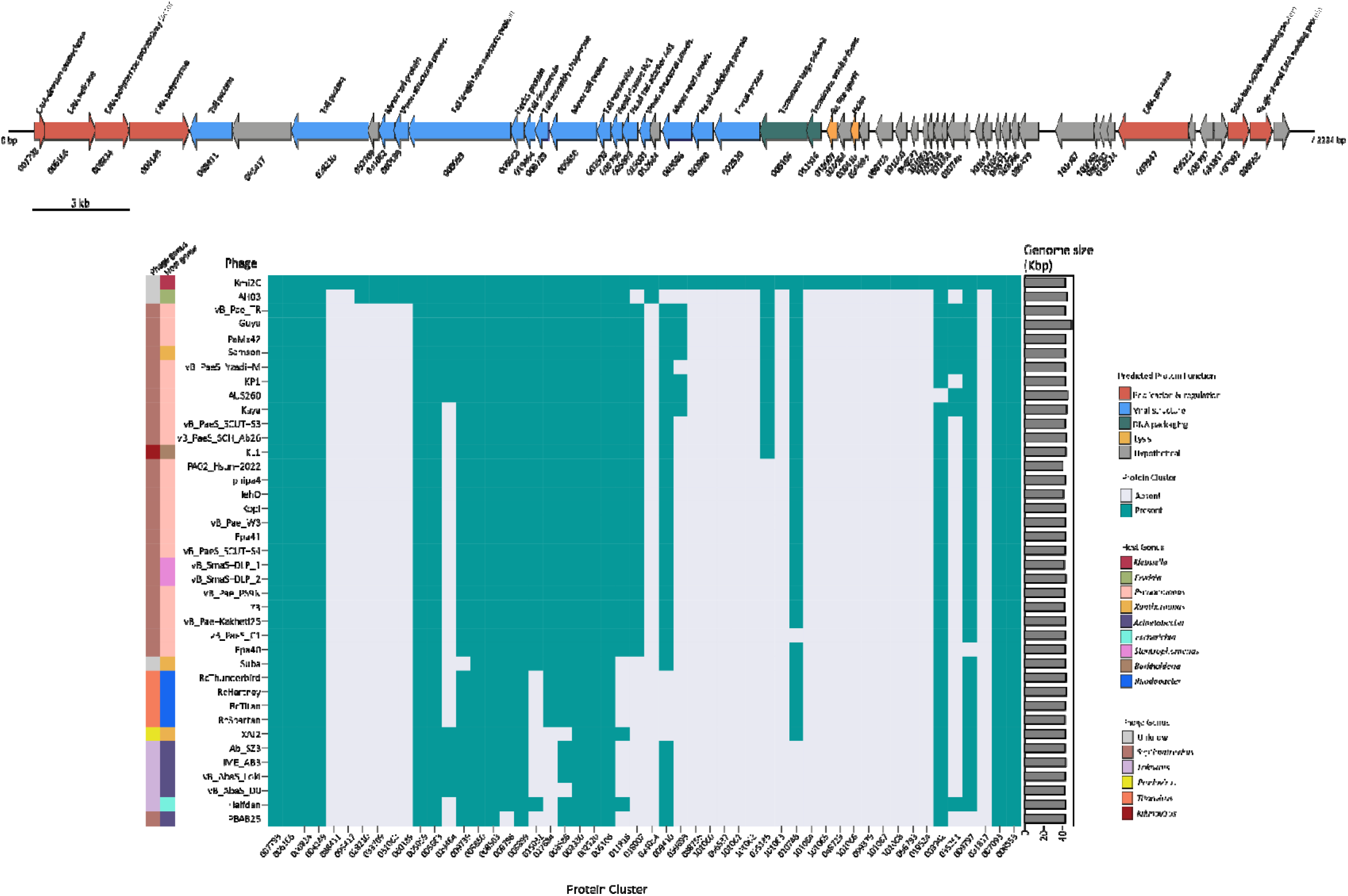
Genome map and predicted function of vB_KmiS-Kmi2C (upper image) and presence/absence of PCs within VC (lower image). The genome map shows the predicted function of the coding regions, direction of transcription and PC ID for each gene segment. The phage genus and host genus are shown to the left of the PC heatmap (see legend for annotation) and phage genome size on the right-hand side of the heatmap.

Analysis using NCBI BLASTN showed the genome of vB_KmiS-Kmi2C shared 85.6 % identity (80 % query coverage) with a phage metagenome-assembled genome (accession OP072809) recovered from human faeces in Japan (Nishijima *et al*., 2022). Preliminary analysis of the genome sequence using the online version of ViPTree suggested the phage was novel (not shown). Further support for this came from genomic similarity analysis (**Supplementary Figure 2**) and analyses of protein–protein network data using vConTACT (**Figure 5**). According to vConTACT, vB_KmiS-Kmi2C shared a VC with 38 other known phage from a range of genera: *Septimatrevius* (n=24), *Lokivirus* (n=6), *Titanvirus* (n=4), *Pradovirus* (n=1), *Kilunavirus* (n=1) and 2 unclassified genera. The VC did not contain any other phage isolated using *Klebsiella* and the majority of phage had *Pseudomonas* as an assigned host (**Figure 6**). The intergenomic similarity within the VC ranged from 100 % to 2 %, with the highest similarity noted between phage within the same genus (**Supplementary Figure 2**). vB_KmiS-Km2iC showed the highest genomic similarity (18.8 %) to *Septimatrevirus* vB_Pae-Kakheti25, a *Pseudomonas* phage. The genera *Titanvirus*, *Lokivirus*, *Septimatrevirus* do not currently belong to any recognised phage family, whereas *Pradovirus* belongs to the *Autographiviridae*, which are podovirus-like in terms of their morphology. Therefore, we suggest vB_KmiS-Kmi2C does not belong to any currently recognised phage family.

**Figure 5.**
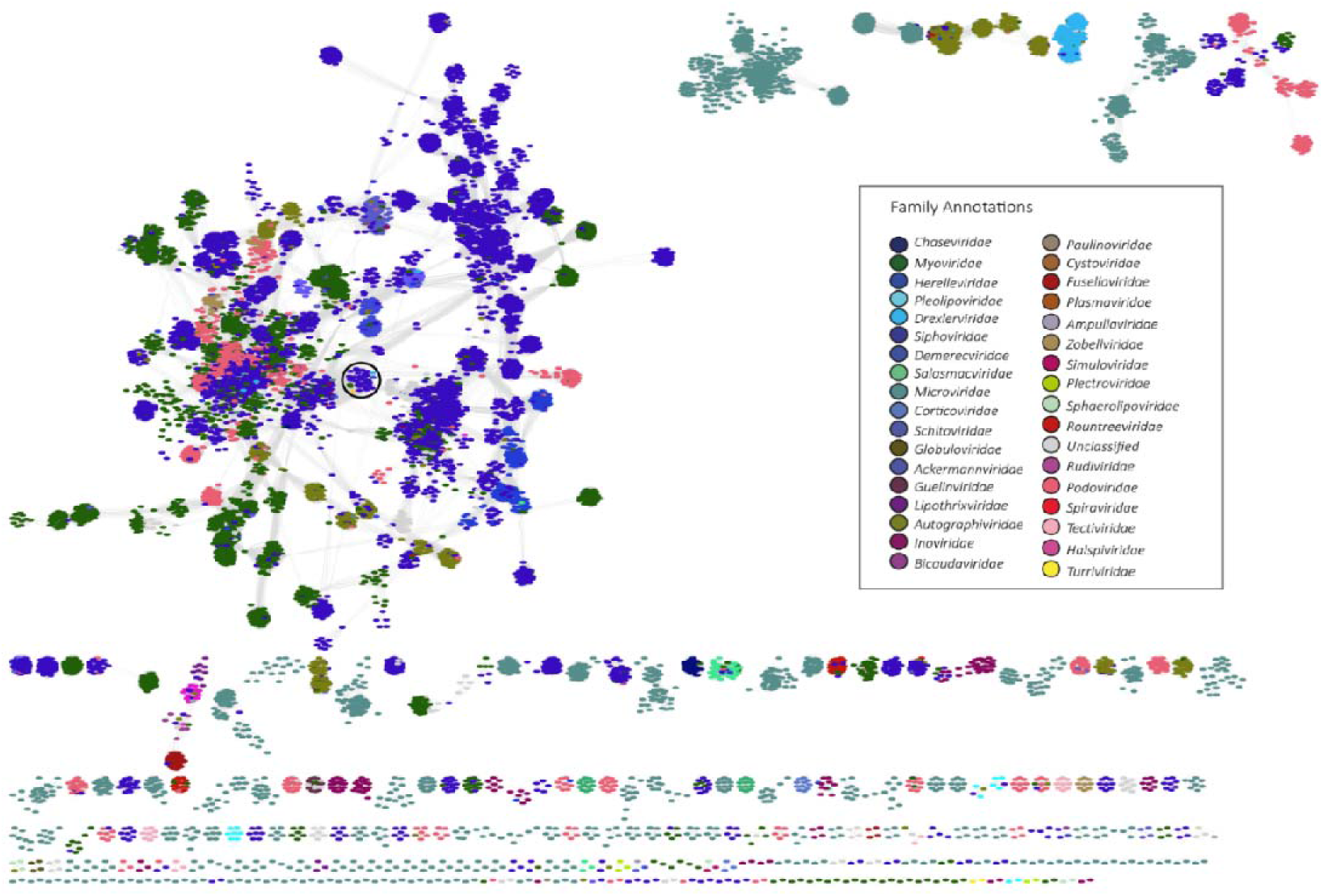
Protein-sharing network generated from vB_KmiS-Kmi2C and INPHARED database. Each node represents a phage genome and connection between the genomes shown by a grey line. The nodes are annotated according to viral family. The VC containing vB_KmiS-Kmi2C is highlighted with a black circle.

**Figure 6.**
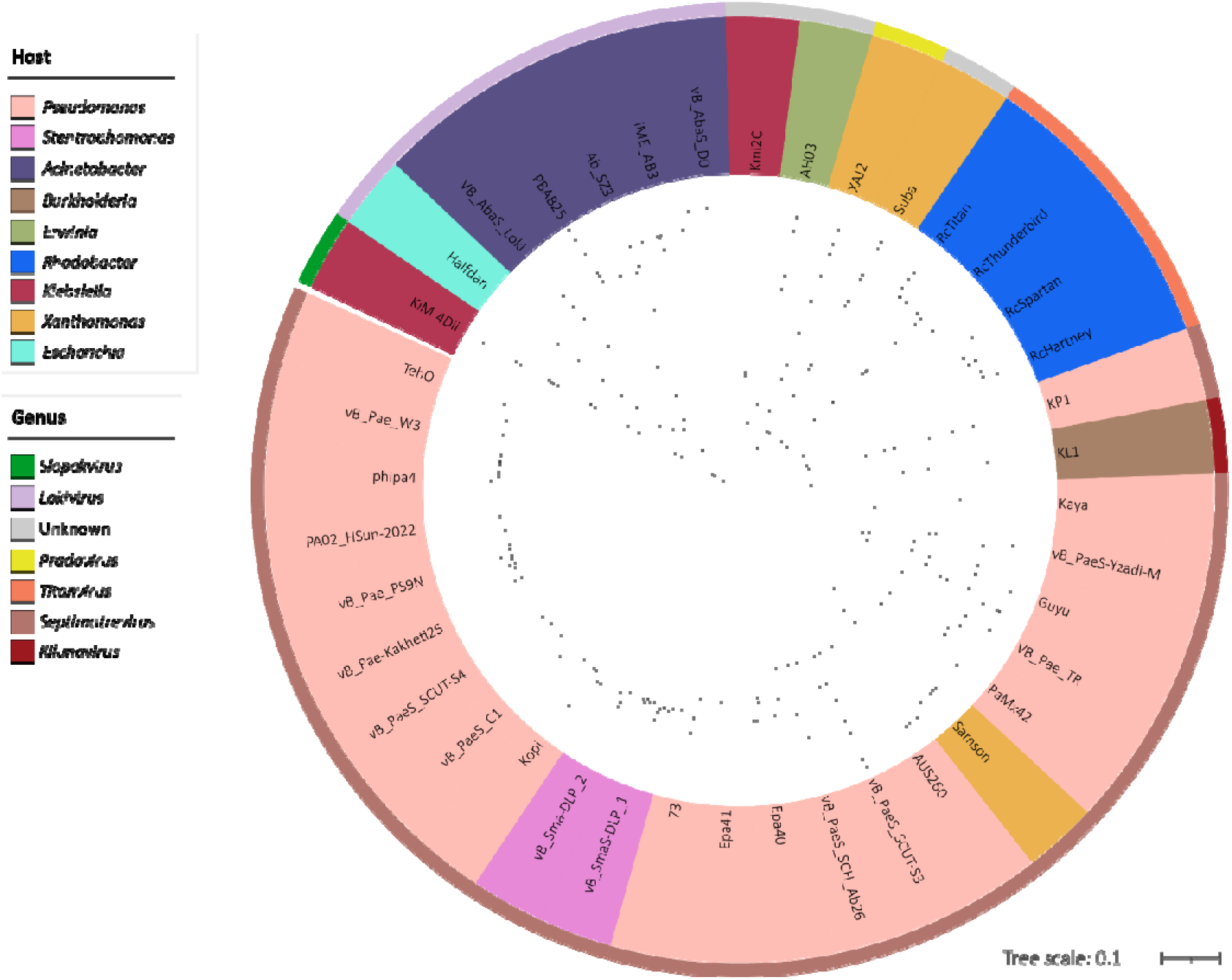
Protein phylogenetic analysis generated with VipTree from all phage within the VC. The inner circle is coloured according to phage host and outer circle according to assigned phage genus. The tree was rooted at the midpoint and scale shown in the bottom right. *Klebsiella* phage vB_KmiM-4Dii (KiM 4Dii), a *Slopekvirus* phage (Smith-Zaitlik *et al*., 2022), was used as an outlier.

The vConTACT output was used to determine the shared protein clusters (PCs) within the VC compared to phage vB_KmiS-Kmi2C (lower panel **Figure 4**). Of the 53 PCs identified in vB_KmiS-Kmi2C, 14 were shared between all phage within the VC (007793 Cas4-domain exonuclease (KJBENDCP_00001); 006166 DNA helicase (KJBENDCP_00002); 006824 DNA polymerase processivity factor (KJBENDCP_00003); 004149 DNA polymerase (KJBENDCP_00004); 005059 tail length tape measure protein (KJBENDCP_00011); 005663 Neck1 protein (KJBENDCP_00012); 005850 minor tail protein (KJBENDCP_00015); 008503 tail terminator (KJBENDCP_00016); 005899 head-tail adaptor Ad1 (KJBENDCP_00018); 003930 head scaffolding protein (KJBENDCP_00022); 002520 portal protein (KJBENDCP_00023); 005106 terminase large subunit (KJBENDCP_00024); 007093 Sak4-like ssDNA annealing protein (KJBENDCP_00053); 008553 ssDNA binding protein (KJBENDCP_000154)). Seven of the shared PCs were associated with viral structure and six were associated with viral replication/regulation. vB_KmiS-Kmi2C shared the most PCs with *Erwinia* phage AH03, including most of the viral structure genome module.

Protein-based phylogenetic analysis with VipTree showed phages within the VC grouped according to the assigned genus, with vB_Kmi2S_Kmi2C not within the defined genus groups (**Figure 6**). vB_KmiS-Kmi2C showed the closest relationship to *Erwinia* phage AH03, as previously noted in the lower panel of **Figure 4**. A representative species from three closely related genera were included in a further protein-based phylogenetic analysis to determine if vB_KmiS-Kmi2C grouped with genera outside the VC defined by vConTACT: *Nipunavirus* NP1 (NC_031058), *Nonagavirus* JenK1 (NC_024146) and *Seuratvirus* Cajan (NC_028776) (**Supplementary Figure 3**). Analysis of the shared protein percentage within the viral cluster was consistent with the intergenomic similarity and protein phylogenetic analysis (**Supplementary Figure 4**). The phage belonging to the same genera shared the highest percentage of proteins. vB_KmiS-Kmi2C shared the highest percentage of proteins with *Pseudomonas* phage *Septimatrevirus* KP1 and *Erwinia* phage AH03. This confirmed that vB_KmiS-Kmi2C does not belong to any currently defined genus of phage.

Given that vB_KmiS-Kmi2C represents a novel family, genus and species of phage, we propose the following taxonomy: *Duplodnaviria* › *Heunggongvirae* › *Uroviricota* › *Caudoviricetes* › *Dilsviridae* › *Dilsvirus* › *Dilsvirus Kmi2C*.

## DISCUSSION

There has been a significant research effort into the investigation of phages for the treatment of *K. pneumoniae* infections and we have recently authored an extensive review on phages of *Klebsiella* spp. (Herridge *et al*., 2020). However, there are relatively few studies documenting the isolation and characterisation of lytic phages against members of the KoC (Kęsik-Szeloch *et al*., 2013; Townsend *et al*., 2021; Smith-Zaitlik *et al*., 2022), an emerging group of clinical and veterinary pathogens with growing antibiotic resistance. As such, we set out to identify phages with the capacity to lyse clinically relevant members of the KoC. We have previously reported on the characterisation of two such phages with lytic activity against numerous species within the KoC (Smith-Zaitlik *et al*., 2022). Here, we report on the characterisation of phage vB_KmiS-Kmi2C isolated against a carbapenem-resistant isolate of *K. michiganensis* (PS_Koxy2) (Shibu *et al*., 2021), for which we generated a hybrid genome assembly and phenotypic data to allow it to be differentiated from closely related strains (PS_Koxy1, PS_Koxy4) from our in-house collection.

There were clear differences in the biofilm-forming abilities of the three GES-5-positive strains, yet all strains encoded the same virulence factors (**Supplementary Table 2**). We did not compare protein sequences of the virulence factors across the strains. It is acknowledged that our understanding of KoC phenotypes is poor (Yang *et al*., 2022), and our future work will look to address how differences in biofilm-forming abilities of KoC members and their genetic differences contribute to survival and potentially pathogenicity.

While the three strains all carried two small (4,448 and 8,300 bp) plasmids, only PS_Koxy2 and PS_Koxy4 carried a 76,860-bp plasmid. This and the other larger plasmids carried by the strains will be described elsewhere as part of ongoing work. The 8,300-bp IncQ plasmid carried by the three strains has been described elsewhere (Ellington *et al*., 2019). While the 4,448-bp plasmid (mobility group MOBP1, replicon type Col440I) has been detected in a strain of *K. michiganensis* previously (AbuOun *et al*., 2021), it has not been described in detail. Based on its gene content (specifically, MobA/MobC), we predict the plasmid to be conjugative plasmid. The MobA and MobC functional homologues MbeA and MbeC, respectively, are required for efficient mobilisation of the ColE1 plasmid (Varsaki *et al*., 2012).

### Host range analysis and anti-biofilm properties

Understanding the lytic profile of any individual phage is important for its potential deployment in a clinical setting, either in an isolated form or for inclusion in a phage cocktail that may increase host range and potentially suppress phage resistance (Yang *et al*., 2020). Ordinarily, phage host range is narrow, sometimes down to the strain level. In accordance with this, we found that vB_KmiS-Kmi2C exhibits a relatively narrow host range, lysing six clinical isolates from the KoC out of a panel of 84 isolates representing a range of clinical and animal *Klebsiella* spp.

To further characterise the therapeutic potential of vB_KmiS-Kmi2C, experiments were performed to determine the phage’s capacity to prevent and disrupt biofilms formed by the clinical isolates susceptible to lysis in our host range analysis. Biofilm formation can reduce the efficacy of antibiotic chemotherapy and contributes to the successful colonisation of wounds and indwelling medical devices (Wang *et al*., 2020). Therefore, phages with anti-biofilm properties may make particularly attractive therapeutics. We found that vB_KmiS-Kmi2C is highly effective at both preventing biofilm formation and disrupting established biofilms. It is likely that prevention of biofilm formation is largely due to the lytic action of the phage, resulting in cell death and the prevention of biofilm formation. However, established biofilms can be extremely recalcitrant to disruption. This is likely due to the complex nature of mature biofilms which are typically composed of numerous macromolecules including, polysaccharides, proteins, nucleic acids and metabolically inactive cells (Chhibber, Nag & Bansal, 2013). Phage-mediated disruption of *K. michiganensis* biofilms has been previously reported in the literature (Ku *et al*., 2021). The lytic phage KMI8 was shown to disrupt a *K*. *michiganensis* mono-biofilm produced by a strain expressing the polysaccharide capsule KL70 locus. It was hypothesised this antibiofilm activity was largely due to the activity of a potential phage-associated depolymerase, identified as a putative endosialidase encoded within the KMI8 genome. The antibiofilm effects of phages against *K. oxytoca* have also been investigated (Townsend, Moat & Jameson, 2020). A recent study showed that phages originally isolated against *K. pneumoniae* were able to reduce the viability of *K. oxytoca* biofilms when added at the start of biofilm formation. However, the same phages were unable to reduce the viability of established *K. oxytoca* biofilms. Here, we found that vB_KmiS-Kmi2C is highly effective at disrupting established biofilms formed by both *K. michiganensis* and *K. oxytoca*. In the present study, we found no evidence of depolymerase activity associated with vB_KmiS-Kmi2C. Plaques on susceptible host lawns did not show evidence of haloes which are typically indicative of depolymerase activity. Furthermore, our bioinformatic analysis did not identify any genes associated with depolymerase activity. Studies are therefore ongoing to determine the mechanism of biofilm disruption used by vB_KmiS-Kmi2C.

### Phage vB_KmiS-Kmi2C represents a new family and genus

Whole-genome sequencing is important for the identification of features relevant to phage therapy and for aiding taxonomic classification. Genomic analysis using the resistance gene identifier tool available through the comprehensive antibiotic resistance database did not identify the presence of known antimicrobial resistance mechanisms within the genome of vB_KmiS-Kmi2C. The presence of such genes would suggest the potential to transfer antibiotic resistance markers between bacteria if vB_KmiS-Kmi2C were to be applied therapeutically in a polymicrobial setting. PhageLeads did not detect the presence of classical lysogeny genes, suggesting vB_KmiS-Kmi2C is exclusively lytic.

Initial genome sequence analysis of vB_KmiS-Kmi2C using BLASTN and ViPTree suggested that our phage was novel, sharing little sequence similarity with other deposited phage genomes. To aid with taxonomy, we employed the gene-sharing network analysis tool vConTACT2. Phage taxonomic assignment using whole genome gene-sharing profiles has been shown to be highly accurate; a recent study showed that vConTACT2 produces near-identical replication of existing genus-level viral taxonomy assignments from the International Committee on Taxonomy of Viruses (ICTV) (Bin Jang *et al*., 2019). Our analysis using vConTACT2 and the INPHRARED database of phage genomes identified that vB_KmiS-Kmi2C shared a VC with 38 other sequenced phages. The VC was polyphyletic, containing phage from at least five different genera, suggesting that vB_KmiS-Km2iC did not group with any existing genus. Recently published guidelines have suggested that for genome-based phage taxonomy, assignment to taxonomic ranks should be based on nucleotide sequence identity across entire genomes (Adriaenssens & Brister, 2017; Turner *et al*., 2021). These guidelines suggest any two phages belong to the same species if they are more than 95 % identical across their genome and belong to the same genus if they are more than 70 % identical. We used VIRIDIC to determine the intergenomic similarity between vB_KmiS-Kmi2C and the 38 other phages identified within the VC. Our findings show that based on these criteria, vB_KmiS-Kmi2C cannot be assigned to any current species or genus as it shares only 18.8 % genomic similarity to *Septimatrevirus* vB_Pae-Kakheti25, a *Pseudomonas* phage, the closest nucleotide sequence relative we could identify.

Most of the phages within the VC (21/38) were identified as infecting bacteria belonging to the genus *Pseudomonas*. According to the GenBank accession details associated with the genomes of these phages, they were all isolated-on strains of *P. aeruginosa*. A preliminary screen of vB_KmiS-Kmi2C against a panel of 11 *P. aeruginosa* isolates did not identify any lytic activity against members of this species (data not shown).

In addition to identifying phages belonging to a VC, vConTACT2 can also identify proteins shared by members within the VC. Our analysis found that 14 proteins were shared between all phages within the VC, of which seven were associated with viral structure and six were associated with viral replication/regulation. At least 35 of the phages within the VC are from genera with siphovirus-like morphology. It is therefore unsurprising that phages within the cluster share several proteins related to general structure/morphology.

Phage vB_KmiS-Kmi2C is the only member of the VC known to infect members of the genus *Klebsiella*. Host-cell tropism is often determined by receptor-binding proteins (RBPs) which mediate recognition and attachment to host cells (Dams *et al*., 2019). RBPs are typically identified as phage tail fibres or spikes (Nobrega *et al*., 2018). The genomic region of vB_KmiS-Kmi2C encoding hypothetical tail fibre proteins can be seen in **Figure 4** and comprises proteins 088411, 095417, 028210, 033709, 031062 and 060189 (KJBENDCP_00005 to KJBENDCP_00010). Unannotated proteins within this region were confirmed to share homology with phage tail fibre proteins by HHPred analysis using an e-value cut-off of >10^-5^. The tail fibre proteins of vB_KmiS-Kmi2C are not widely present within the VC and are typically shared by only one or two phages within the cluster. This may explain the host tropism of vB_KmiS-Kmi2C compared to other members of the VC but further experimentation will be required to determine this. Protein 005059 (KJBENDCP_00011) is a predicted tail length tape measure protein; such proteins are involved in determining tail length and participate in DNA injection into the host-cells and are not involved in host recognition (Mahony *et al*., 2016).

Our additional protein-based phylogenetic analyses, including phages from outside the VC, highlighted that vB_Kmi2S_Kmi2C does not group with any genera included in the analyses and as such does not belong to any currently defined genus of phage.

Phages are recognised as the most abundant biological entities on the planet and exhibit extensive biological diversity (Hendrix, 2002). With an estimated 10^31^ phage virions in the biosphere (Hatfull, 2015) and 14,244 complete phage genomes sequenced as of 2021 (Cook *et al*., 2021), it is clear we have characterised a relatively small number of distinct phages. However, with the cost of genome sequencing falling and the capacity to assemble phage genomes from metagenome sequence data, it is likely that the number of novel phages identified will increase rapidly. The classification of phages representing entirely novel genera or families/sub-families poses a challenge for taxonomy, especially when new isolates share little sequence similarity with previously characterised phages. Previously, tailed phages were assigned to a family based on morphology, with the families *Podoviridae*, *Myoviridae* and *Siphoviridae* belonging to the order *Caudovirales*.

However, the increase in available phage sequence data has shown that these families are paraphyletic. It has, therefore, been suggested that they are abolished, and that phage morphology should no longer play a role in classification (Turner *et al*., 2021). Based on recently updated guidelines for the classification of phage based on nucleotide sequence identity, and subsequent protein based phylogenetic analyses, we show here that phage vB_KmiS-Kmi2C represents a new genus.

## AUTHOR CONTRIBUTION STATEMENT

Conceptualisation: FN, PS, LH, DN. Data curation: all authors. Formal analysis: FN, TSZ, PS, ME, ALM, LH, DN. Funding acquisition: PS, ALM, LH. Investigation: FN, TSZ, PS, ME, ALM, LH, DN. Methodology: FN, ME, LH, DN. Resources: ALM, LH, DN. Supervision: ALM, LH, DN. Visualisation: FN, LH, DN. Writing – original draft: FN, ME, LH, DN. Writing – reviewing and editing: all authors.

## ACKNOWLEDGEMENTS

PS was in receipt of an IBMS Research Grant (project title “Isolation of lytic bacteriophages active against antibiotic-resistant *Klebsiella pneumoniae*”). Imperial Health Charity is thanked for contributing to registration fees for the Professional Doctorate studies of PS. ME is supported by the Egyptian Government – Ministry of Higher Education/Egyptian Cultural and Education Bureau. This work used computing resources funded by the Research Contingency Fund of the Department of Biosciences, Nottingham Trent University (NTU). TSZ completed this work as part of an MRes degree at NTU.

## Abbreviations

CARD: comprehensive antibiotic resistance database
KoC: *Klebsiella oxytoca* complex
PC: protein cluster
RBP: receptor binding protein
SD: standard deviation
TEM: transmission electron microscopy
TSA: tryptone soy agar
TSBG: tryptone soy broth supplemented with 1 % glucose
VC: viral cluster
VFDB: virulence factor database

## REFERENCES

AbuOun, M., Jones, H., Stubberfield, E., Gilson, D., Shaw, L.P., Hubbard, A.T.M., Chau, K.K., Sebra, R., Peto, T.E.A., Crook, D.W., Read, D.S., Gweon, H.S., Walker, A.S., Stoesser, N., Smith, R.P., Anjum, M.F. and On Behalf Of The Rehab Consortium, null (2021) A genomic epidemiological study shows that prevalence of antimicrobial resistance in Enterobacterales is associated with the livestock host, as well as antimicrobial usage. Microb Genomics. 7, 000630.

Adriaenssens, E. and Brister, J.R. (2017) How to Name and Classify Your Phage: An Informal Guide. Viruses. 9, E70.

Alcock, B.P., Raphenya, A.R., Lau, T.T.Y., Tsang, K.K., Bouchard, M., Edalatmand, A., Huynh, W., Nguyen, A.-L.V., Cheng, A.A., Liu, S., Min, S.Y., Miroshnichenko, A., Tran, H.-K., Werfalli, R.E., Nasir, J.A., Oloni, M., Speicher, D.J., Florescu, A., Singh, B., Faltyn, M., Hernandez-Koutoucheva, A., Sharma, A.N., Bordeleau, E., Pawlowski, A.C., Zubyk, H.L., Dooley, D., Griffiths, E., Maguire, F., Winsor, G.L., Beiko, R.G., Brinkman, F.S.L., Hsiao, W.W.L., Domselaar, G.V. and McArthur, A.G. (2020) CARD 2020: antibiotic resistome surveillance with the comprehensive antibiotic resistance database. Nucleic Acids Res. 48, D517–D525.

Bankevich, A., Nurk, S., Antipov, D., Gurevich, A.A., Dvorkin, M., Kulikov, A.S., Lesin, V.M., Nikolenko, S.I., Pham, S., Prjibelski, A.D., Pyshkin, A.V., Sirotkin, A.V., Vyahhi, N., Tesler, G., Alekseyev, M.A. and Pevzner, P.A. (2012) SPAdes: a new genome assembly algorithm and its applications to single-cell sequencing. J Comput Biol J Comput Mol Cell Bio.l19, 455–477.

Bin Jang, H., Bolduc, B., Zablocki, O., Kuhn, J.H., Roux, S., Adriaenssens, E.M., Brister, J.R., Kropinski, A.M., Krupovic, M., Lavigne, R., Turner, D. and Sullivan, M.B. (2019) Taxonomic assignment of uncultivated prokaryotic virus genomes is enabled by gene-sharing networks. Nat Biotechnol. 37, 632–639.

Bolduc, B., Jang, H.B., Doulcier, G., You, Z.-Q., Roux, S. and Sullivan, M.B. (2017) vConTACT: an iVirus tool to classify double-stranded DNA viruses that infect Archaea and Bacteria. PeerJ. 5, e3243.

Bolger, A.M., Lohse, M. and Usadel, B. (2014) Trimmomatic: a flexible trimmer for Illumina sequence data. Bioinforma Oxf Engl. 30, 2114–2120.

Chhibber, S., Nag, D. and Bansal, S. (2013) Inhibiting biofilm formation by Klebsiella pneumoniae B5055 using an iron antagonizing molecule and a bacteriophage. BMC Microbiol. 13, 174.

Clokie, M.R., Millard, A.D., Letarov, A.V. and Heaphy, S. (2011) Phages in nature. Bacteriophage. 1, 31–45.

Cook, R., Brown, N., Redgwell, T., Rihtman, B., Barnes, M., Clokie, M., Stekel, D.J., Hobman, J., Jones, M.A. and Millard, A. (2021) INfrastructure for a PHAge REference Database: Identification of Large-Scale Biases in the Current Collection of Cultured Phage Genomes. PHAGE. 2, 214–223.

Cosic, A., Leitner, E., Petternel, C., Galler, H., Reinthaler, F.F., Herzog-Obereder, K.A., Tatscher, E., Raffl, S., Feierl, G., Högenauer, C., Zechner, E.L. and Kienesberger, S. (2021) Variation in Accessory Genes Within the*Klebsiella oxytoca* Species Complex Delineates Monophyletic Members and Simplifies Coherent Genotyping. Front Microbiol. 12, 692453.

Dams, D., Brøndsted, L., Drulis-Kawa, Z. and Briers, Y. (2019) Engineering of receptor-binding proteins in bacteriophages and phage tail-like bacteriocins. Biochem Soc Trans. 47, 449–460.

Eladawy, M., El-Mowafy, M., El-Sokkary, M.M.A. and Barwa, R. (2021) Antimicrobial resistance and virulence characteristics in ERIC-PCR typed biofilm forming isolates of P. aeruginosa. Microb Pathog. 158, 105042.

Ellington, M.J., Davies, F., Jauneikaite, E., Hopkins, K.L., Turton, J.F., Adams, G., Pavlu, J., Innes, A.J., Eades, C., Brannigan, E.T., Findlay, J., White, L., Bolt, F., Kadhani, T., Chow, Y., Patel, B., Mookerjee, S., Otter, J.A., Sriskandan, S., Woodford, N. and Holmes, A. (2019) A multi-species cluster of GES-5 carbapenemase producing Enterobacterales linked by a geographically disseminated plasmid. Clin Infect Dis Off Publ Infect Dis Soc Am .

Giménez, M., Ferrés, I. and Iraola, G. (2022) Improved detection and classification of plasmids from circularized and fragmented assemblies. bioRxiv.

Gu, Z., Eils, R. and Schlesner, M. (2016) Complex heatmaps reveal patterns and correlations in multidimensional genomic data. Bioinforma Oxf Engl . 32, 2847–2849.

Guy, L., Kultima, J.R. and Andersson, S.G.E. (2010) genoPlotR: comparative gene and genome visualization in R. Bioinforma Oxf Engl. 26, 2334–2335.

Haines, M.E.K., Hodges, F.E., Nale, J.Y., Mahony, J., van Sinderen, D., Kaczorowska, J., Alrashid, B., Akter, M., Brown, N., Sauvageau, D., Sicheritz-Pontén, T., Thanki, A.M., Millard, A.D., Galyov, E.E. and Clokie, M.R.J. (2021) Analysis of Selection Methods to Develop Novel Phage Therapy Cocktails Against Antimicrobial Resistant Clinical Isolates of Bacteria. Front Microbiol. 12, 613529.

Hatfull, G.F. (2015) Dark Matter of the Biosphere: the Amazing World of Bacteriophage Diversity. J Virol. 89, 8107–8110.

Hendrix, R.W. (2002) Bacteriophages: Evolution of the Majority. Theor Popul Biol. 61, 471–480.

d’Herelle, F. (1918) Sur le rôle du microbe filtrant bactériophage dans la dysentérie bacillaire. Comptes Rendus Académie Sci . 167, 970–972.

Herridge, W.P., Shibu, P., O’Shea, J., Brook, T.C. and Hoyles, L. (2020) Bacteriophages of Klebsiella spp., their diversity and potential therapeutic uses. J Med Microbiol. 69, 176–194.

Kalyaanamoorthy, S., Minh, B.Q., Wong, T.K.F., von Haeseler, A. and Jermiin, L.S. (2017) ModelFinder: fast model selection for accurate phylogenetic estimates. Nat Methods. 14, 587–589.

Kęsik-Szeloch, A., Drulis-Kawa, Z., Weber-Dąbrowska, B., Kassner, J., Majkowska-Skrobek, G., Augustyniak, D., Łusiak-Szelachowska, M., Żaczek, M., Górski, A. and Kropinski, A.M. (2013) Characterising the biology of novel lytic bacteriophages infecting multidrug resistant *Klebsiella pneumoniae*. Virol J. 10, 100.

Ku, H., Kabwe, M., Chan, H.T., Stanton, C., Petrovski, S., Batinovic, S. and Tucci, J. (2021) Novel Drexlerviridae bacteriophage KMI8 with specific lytic activity against Klebsiella michiganensis and its biofilms. PLOS ONE. 16, e0257102.

Lee, S.H., Ruan, S.-Y., Pan, S.-C., Lee, T.-F., Chien, J.-Y. and Hsueh, P.-R. (2019) Performance of a multiplex PCR pneumonia panel for the identification of respiratory pathogens and the main determinants of resistance from the lower respiratory tract specimens of adult patients in intensive care units. J Microbiol Immunol Infect. 52, 920–928.

Liu, B., Zheng, D., Jin, Q., Chen, L. and Yang, J. (2019) VFDB 2019: a comparative pathogenomic platform with an interactive web interface. Nucleic Acids Res. 47, D687–D692.

Lowe, C., Willey, B., O’Shaughnessy, A., Lee, W., Lum, M., Pike, K., Larocque, C., Dedier, H., Dales, L., Moore, C., McGeer, A., and the Mount Sinai Hospital Infection Control Team (2012) Outbreak of Extended-Spectrum β-Lactamase‒producing *Klebsiella oxytoca* Infections Associated with Contaminated Handwashing Sinks1. Emerg Infect Dis. 18, 1242–1247.

Mahony, J., Alqarni, M., Stockdale, S., Spinelli, S., Feyereisen, M., Cambillau, C. and Sinderen, D. van (2016) Functional and structural dissection of the tape measure protein of lactococcal phage TP901-1. Sci Rep. 6, 36667.

Merla, C., Rodrigues, C., Passet, V., Corbella, M., Thorpe, H.A., Kallonen, T.V.S., Zong, Z., Marone, P., Bandi, C., Sassera, D., Corander, J., Feil, E.J. and Brisse, S. (2019) Description of Klebsiella spallanzanii sp. nov. and of Klebsiella pasteurii sp. nov. Front Microbiol. 10, 2360.

Merritt, J.H., Kadouri, D.E. and O’Toole, G.A. (2005) Growing and analyzing static biofilms. Curr Protoc Microbiol. Chapter 1, Unit 1B.1.

Minh, B.Q., Nguyen, M.A.T. and von Haeseler, A. (2013) Ultrafast Approximation for Phylogenetic Bootstrap. Mol Biol Evol. 30, 1188–1195.

Moraru, C., Varsani, A. and Kropinski, A.M. (2020) VIRIDIC—A Novel Tool to Calculate the Intergenomic Similarities of Prokaryote-Infecting Viruses. Viruses. 12, 1268.

Nayfach, S., Camargo, A.P., Schulz, F., Eloe-Fadrosh, E., Roux, S. and Kyrpides, N.C. (2021) CheckV assesses the quality and completeness of metagenome-assembled viral genomes. Nat Biotechnol. 39, 578–585.

Nishijima, S., Nagata, N., Kiguchi, Y., Kojima, Y., Miyoshi-Akiyama, T., Kimura, M., Ohsugi, M., Ueki, K., Oka, S., Mizokami, M., Itoi, T., Kawai, T., Uemura, N. and Hattori, M. (2022) Extensive gut virome variation and its associations with host and environmental factors in a population-level cohort. Nat Commun . 13, 5252.

Nishimura, Y., Yoshida, T., Kuronishi, M., Uehara, H., Ogata, H. and Goto, S. (2017) ViPTree: the viral proteomic tree server.Bioinforma Oxf Engl. 33, 2379–2380.

Nobrega, F.L., Vlot, M., de Jonge, P.A., Dreesens, L.L., Beaumont, H.J.E., Lavigne, R., Dutilh, B.E. and Brouns, S.J.J. (2018) Targeting mechanisms of tailed bacteriophages. Nat Rev Microbiol . 16, 760–773.

Paasch, C., Wilczek, S. and Strik, M.W. (2017) Liver abscess and sepsis caused by *Clostridium perfringens* and *Klebsiella oxytoca*. Int J Surg Case Rep. 41, 180–183.

Redondo-Salvo, S., Bartomeus-Peñalver, R., Vielva, L., Tagg, K.A., Webb, H.E., Fernández-López, R. and de la Cruz, F. (2021) COPLA, a taxonomic classifier of plasmids. BMC Bioinformatics. 22, 390.

Schwengers, O., Jelonek, L., Dieckmann, M.A., Beyvers, S., Blom, J. and Goesmann, A. (2021) Bakta: rapid and standardized annotation of bacterial genomes via alignment-free sequence identification. Microb Genomics. 7.

Shakya, P., Shrestha, D., Maharjan, E., Sharma, V.K. and Paudyal, R. (2017) ESBL Production Among *E. coli* and *Klebsiella* spp. Causing Urinary Tract Infection: A Hospital Based Study. Open Microbiol J. 11, 23–30.

Shibu, P., McCuaig, F., McCartney, A.L., Kujawska, M., Hall, L.J. and Hoyles, L. (2021) Improved molecular characterization of the *Klebsiella oxytoca* complex reveals the prevalence of the kleboxymycin biosynthetic gene cluster. Microb Genomics. 7, 000592.

Smith-Zaitlik, T., Shibu, P., McCartney, A.L., Foster, G., Hoyles, L. and Negus, D. (2022) Extended genomic analyses of the broad-host-range phages vB_KmiM-2Di and vB_KmiM-4Dii reveal slopekviruses have highly conserved genomes. Microbiology. 168, 001247.

Sohn, W.-I., Seo, B.F. and Jung, S.-N. (2012) Forehead Abscess Caused by *Klebsiella oxytoca* With Undiagnosed Type 2 Diabetes: J Craniofac Surg. 23, 247–249.

Stepanovic, S., Vukovic, D., Dakic, I., Savic, B. and Svabic-Vlahovic, M. (2000) A modified microtiter-plate test for quantification of staphylococcal biofilm formation. J Microbiol Methods. 40, 175–179.

Taha, O.A., Connerton, P.L., Connerton, I.F. and El-Shibiny, A. (2018) Bacteriophage ZCKP1: A Potential Treatment for Klebsiella pneumoniae Isolated From Diabetic Foot Patients. Front Microbiol. 9, 2127.

Terzian, P., Olo Ndela, E., Galiez, C., Lossouarn, J., Pérez Bucio, R.E., Mom, R., Toussaint, A., Petit, M.-A. and Enault, F. (2021) PHROG: families of prokaryotic virus proteins clustered using remote homology. NAR Genomics Bioinforma. 3, lqab067.

Thompson, J.D., Higgins, D.G. and Gibson, T.J. (1994) CLUSTAL W: improving the sensitivity of progressive multiple sequence alignment through sequence weighting, position-specific gap penalties and weight matrix choice. Nucleic Acids Res. 22, 4673–4680.

Townsend, E.M., Kelly, L., Gannon, L., Muscatt, G., Dunstan, R., Michniewski, S., Sapkota, H., Kiljunen, S.J., Kolsi, A., Skurnik, M., Lithgow, T., Millard, A.D. and Jameson, E. (2021) Isolation and Characterization of *Klebsiella* Phages for Phage Therapy. PHAGE. 2, 26–42.

Townsend, E.M., Moat, J. and Jameson, E. (2020) CAUTI’s next top model - Model dependent Klebsiella biofilm inhibition by bacteriophages and antimicrobials. Biofilm. 2, 100038.

Turner, D., Kropinski, A.M. and Adriaenssens, E.M. (2021) A Roadmap for Genome-Based Phage Taxonomy. Viruses. 13, 506.

Varsaki, A., Lamb, H.K., Eleftheriadou, O., Vandera, E., Thompson, P., Moncalián, G., de la Cruz, F., Hawkins, A.R. and Drainas, C. (2012) Interaction between relaxase MbeA and accessory protein MbeC of the conjugally mobilizable plasmid ColE1. FEBS Lett. 586, 675–679.

Wang, G., Zhao, G., Chao, X., Xie, L. and Wang, H. (2020) The Characteristic of Virulence, Biofilm and Antibiotic Resistance of Klebsiella pneumoniae. Int J Environ Res Public Health. 17, 6278.

Webb, H.E., Kim, J.Y., Tagg, K.A., de la Cruz, F., Peñil-Celis, A., Tolar, B., Ellison, Z., Schwensohn, C., Brandenburg, J., Nichols, M. and Folster, J.P. (n.d.) Genome Sequences of 18 Salmonella enterica Serotype Hadar Strains Collected from Patients in the United States. Microbiol Resour Announc. 11, e00522–22.

White, L., Hopkins, K.L., Meunier, D., Perry, C.L., Pike, R., Wilkinson, P., Pickup, R.W., Cheesbrough, J. and Woodford, N. (2016) Carbapenemase-producing *Enterobacteriaceae* in hospital wastewater: a reservoir that may be unrelated to clinical isolates. J Hosp Infect. 93, 145–151.

Wick, R.R., Judd, L.M., Gorrie, C.L. and Holt, K.E. (2017) Unicycler: Resolving bacterial genome assemblies from short and long sequencing reads. PLoS Comput Biol. 13, e1005595.

Wick, R.R., Schultz, M.B., Zobel, J. and Holt, K.E. (2015) Bandage: interactive visualization of de novo genome assemblies. Bioinforma Oxf Engl. 31, 3350–3352.

Yachison, C.A., Yoshida, C., Robertson, J., Nash, J.H.E., Kruczkiewicz, P., Taboada, E.N., Walker, M., Reimer, A., Christianson, S., Nichani, A., PulseNet Canada Steering Committee and Nadon, C. (2017) The Validation and Implications of Using Whole Genome Sequencing as a Replacement for Traditional Serotyping for a National Salmonella Reference Laboratory. Front Microbiol. 8, 1044.

Yang, J., Long, H., Hu, Y., Feng, Y., McNally, A. and Zong, Z. (2022) Klebsiella oxytoca Complex: Update on Taxonomy, Antimicrobial Resistance, and Virulence. Clin Microbiol Rev. 35, e0000621.

Yang, Y., Shen, W., Zhong, Q., Chen, Q., He, X., Baker, J.L., Xiong, K., Jin, X., Wang, J., Hu, F. and Le, S. (2020) Development of a Bacteriophage Cocktail to Constrain the Emergence of Phage-Resistant *Pseudomonas aeruginosa*. Front Microbiol. 11, 327.

Yukgehnaish, K., Rajandas, H., Parimannan, S., Manickam, R., Marimuthu, K., Petersen, B., Clokie, M.R.J., Millard, A. and Sicheritz-Pontén, T. (2022) PhageLeads: Rapid Assessment of Phage Therapeutic Suitability Using an Ensemble Machine Learning Approach. Viruses. 14, 342.

Zhang, Y. (2008) I-TASSER server for protein 3D structure prediction. BMC Bioinformatics. 9, 40.

Zimmermann, L., Stephens, A., Nam, S.-Z., Rau, D., Kübler, J., Lozajic, M., Gabler, F., Söding, J., Lupas, A.N. and Alva, V. (2018) A Completely Reimplemented MPI Bioinformatics Toolkit with a New HHpred Server at its Core. J Mol Biol. 430, 2237–2243.

