## Supplementary Tables and Figures for "Lytic bacteriophage vB_KmiS-Kmi2C disrupts biofilms formed by members of the *Klebsiella oxytoca* complex, and represents a novel virus family and genus"

**Supplementary Table 1.** Comparison of biofilm-forming abilities of strains with the negative control Ko14

*P* values were calculated using unpaired *t* tests of each biofilm-forming strain against the negative control (Ko14). *P* values were adjusted to account for multiple testing using the Benjamini–Hochberg (BH) method. Mean data are from four technical replicates per biological (Expt) replicate.

|  | **Mean** | | | **SD** | ***P* value** | **BH *P* value** |
| --- | --- | --- | --- | --- | --- | --- |
|  | **Expt 1** | **Expt 2** | **Expt 3** |  |  |  |
| *K. michiganensis* Ko14 | 0.068 | 0.073 | 0.074 | 0.004 |  |  |
| *K. michiganensis* Ko13 | 0.110 | 0.154 | 0.113 | 0.025 | 3.21E-06 | 2.89E-05 |
| *K.* michiganensis Ko43 | 0.169 | 0.198 | 0.206 | 0.020 | 8.77E-06 | 3.95E-05 |
| *K. michiganensis* PS_Koxy4 | 0.238 | 0.276 | 0.295 | 0.029 | 3.14E-04 | 9.43E-04 |
| *K. michiganensis* PS_Koxy2 | 0.156 | 0.158 | 0.160 | 0.002 | 4.86E-04 | 1.09E-03 |
| *K. oxytoca* Ko37 | 0.108 | 0.116 | 0.103 | 0.007 | 8.28E-04 | 1.49E-03 |
| *K. michiganensis* Ko21 | 0.107 | 0.136 | 0.104 | 0.017 | 1.13E-03 | 1.69E-03 |
| *K. oxytoca* Ko53 | 0.178 | 0.225 | 0.194 | 0.024 | 1.06E-02 | 1.36E-02 |
| *K. michiganensis* Ko22 | 0.296 | 0.323 | 0.315 | 0.014 | 1.31E-02 | 1.47E-02 |
| *K. michiganensis* PS_Koxy1 | 0.186 | 0.319 | 0.322 | 0.078 | 2.06E-02 | 2.06E-02 |

**Supplementary Table 2.** Summary of VFDB data for *K. michiganensis* strains PS_Koxy1, PS_Koxy2 and PS_Koxy4

Available as an Excel file (**Supplementary Table 2**).

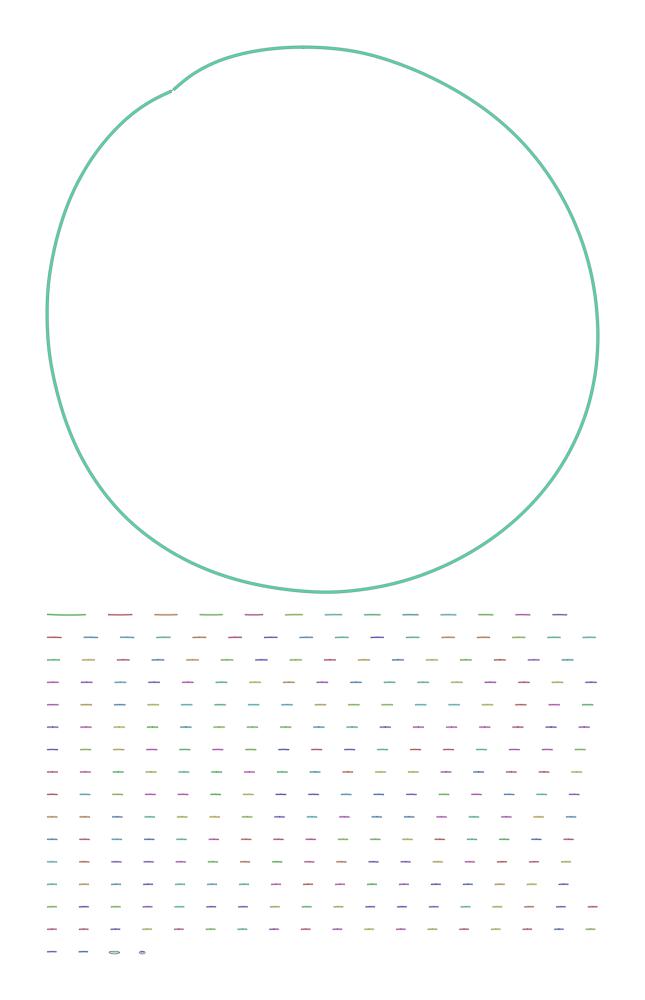

**Supplementary Figure 1.** Bandage image generated from SPAdes assembly of the phage vB_KmiS-Kmi2C genome. The complete circular genome represents the phage, whereas the smaller linear contigs represent host DNA.

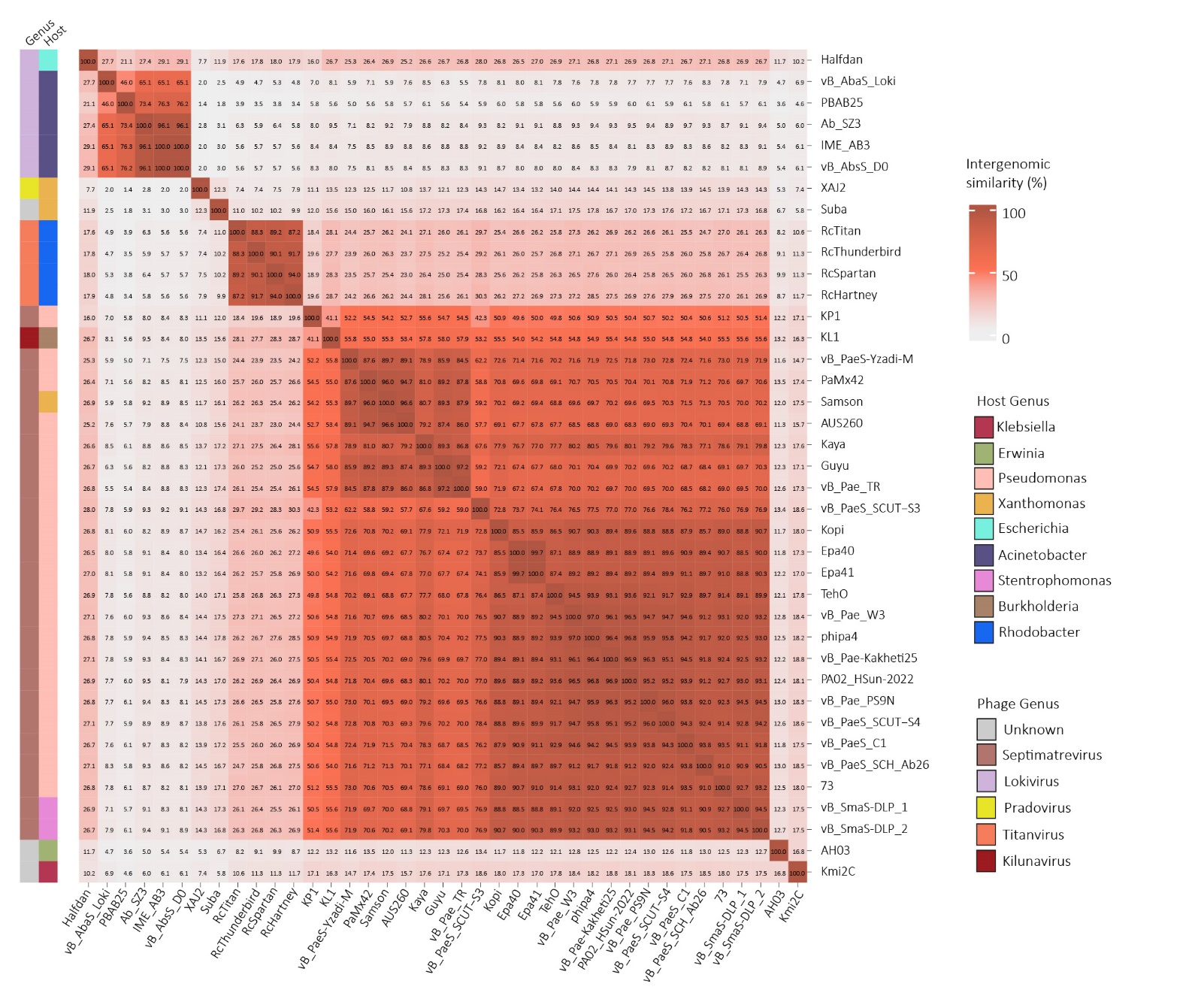

**Supplementary Figure 2.** Intergenomic similarity of all phage within the viral cluster shown by a red gradient (darker red = higher similarity) and percentage within each square. The phage and host genus are represented to the left of the heatmap and coloured annotates in the legend.

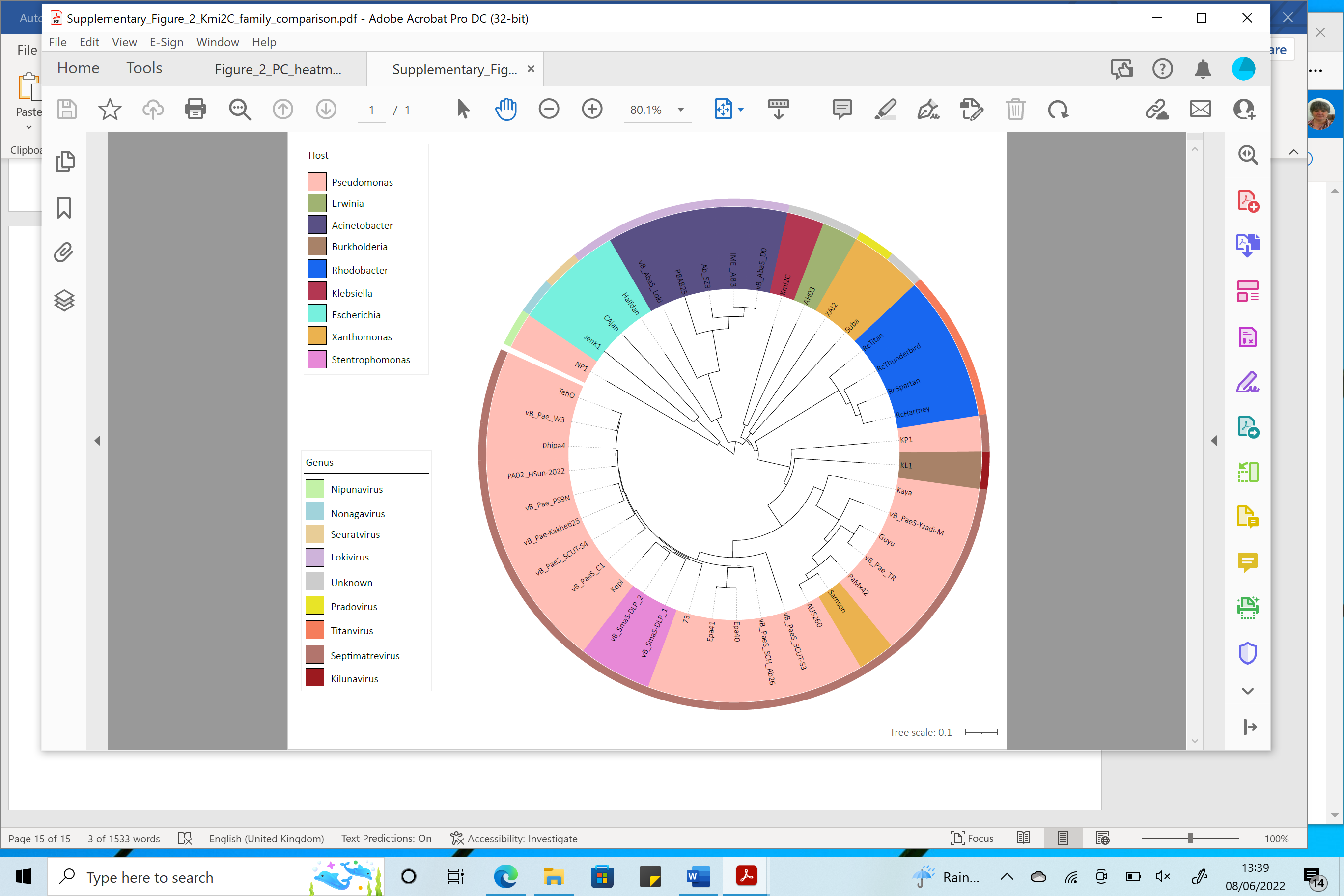

**Supplementary Figure 3.** Protein phylogenetic analysis generated with VipTree from all phage within the viral cluster and 3 representative phage from related genera (*Nipunavirus, Nonagavirus, Seuratvirus*). The inner circle is coloured according to assigned viral genus and outer circle according to phage host. The tree was rooted at the midpoint and scale shown in the bottom right.

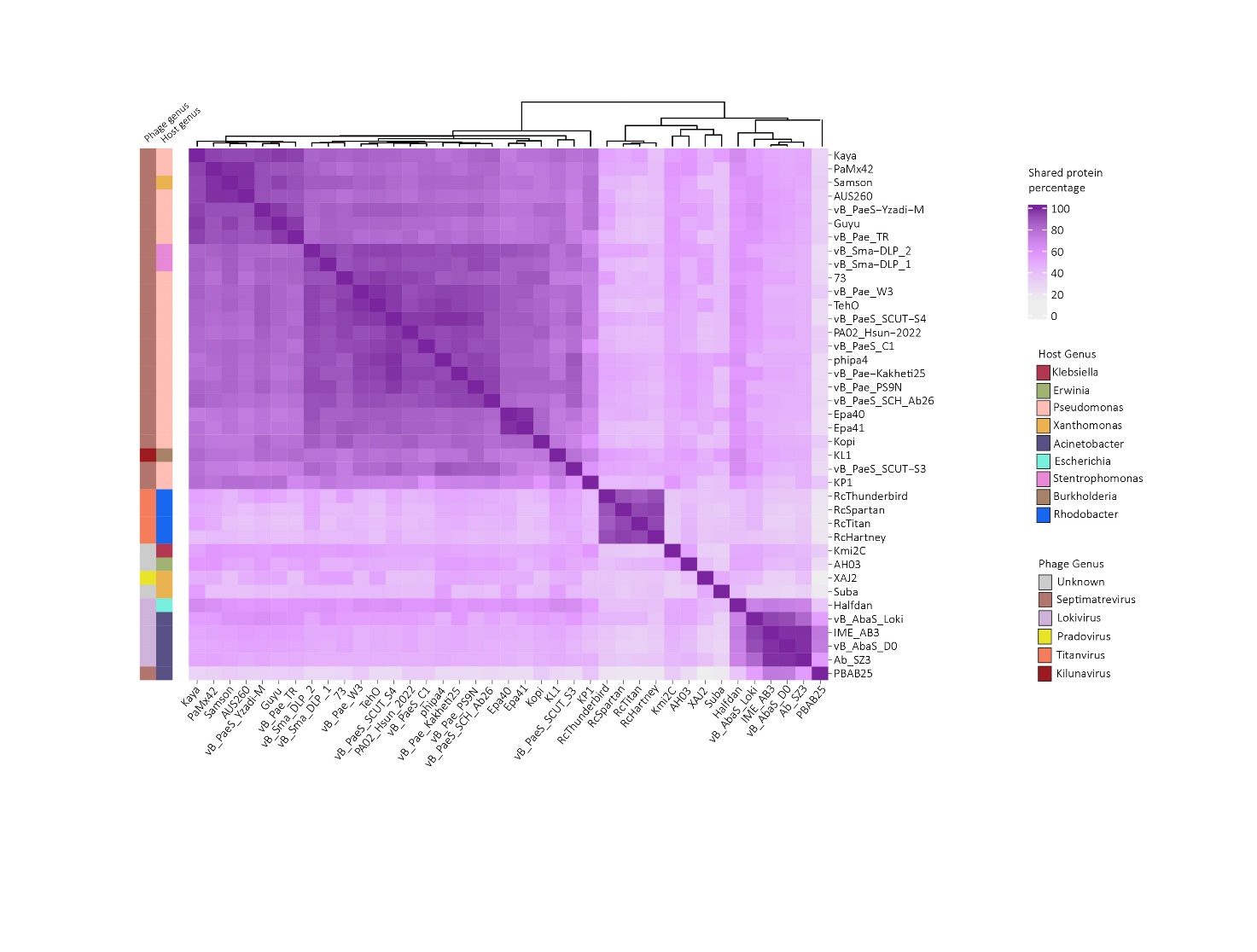

**Supplementary Figure 4.** Shared protein percentage of all phage within the viral cluster shown by a purple gradient (darker purple = higher protein percentage shared). The phage and host genus are represented to the left of the heatmap and coloured annotates in the legend. Dendrogram shown along the top *x* axis.
